# mRNA-GPT: A Generative Model for Full-Length mRNA Design and Optimization

**DOI:** 10.64898/2026.03.31.715707

**Authors:** Sizhen Li, Paul Chauvin, Ofek Gross, Michael Bailey, Sven Jager

## Abstract

We introduce mRNA-GPT, a generative model for end-to-end full-length mRNA sequence design and optimization. Unlike existing approaches that optimize isolated regions, mRNA-GPT jointly optimizes across all three regions (5^′^ UTR, CDS, and 3^′^ UTR) to capture long-range sequence dependencies and cross-region regulatory interactions critical for therapeutic efficacy. The model is pre-trained on 30 million full-length natural mRNA sequences across diverse species and organisms, establishing a robust foundation for sequence generation. We employ Reinforcement Learning (RL), specifically Proximal Policy Optimization (PPO) with oracle-based reward signals, to directly and iteratively optimize target properties, such as half-life and translation efficiency. mRNA-GPT supports flexible generation modes: single regions (UTR or CDS alone), full-length sequences, or generation of any region conditioned on any other region. Through multi-objective optimization, mRNA-GPT achieves Pareto-optimal designs, i.e., solutions for which no single objective can be improved without degrading another, that balance competing properties without sacrificing performance on either objective. mRNA-GPT demonstrates superior design capabilities compared to state-of-the-art methods, achieving enhanced performance in 3^′^ UTR stability optimization, CDS translation rate enhancement, and comprehensive full-length sequence design.

## Introduction

Messenger RNA (mRNA) medicines have transitioned from conceptual promise to clinical reality with the success of COVID-19 vaccines, establishing nucleic-acid delivery as a scalable therapeutic modality [Polack et al., 2020, Baden et al., 2021]. However, therapeutic performance remains exquisitely sensitive to sequence-level determinants distributed across the entire transcript. Coding sequences (CDS) dictate translation kinetics and *in vivo* stability through codon usage and decoding dynamics [Presnyak et al., 2015, Wu et al., 2019]; the 5′ untranslated region (UTR) modulates ribosomal initiation via structural motifs and upstream open reading frames [Leppek et al., 2018, Karollus et al., 2021]; and the 3^′^ UTR integrates microRNA and RNA-binding-protein interactions that collectively govern stability, localization, and translational efficiency [Mayr, 2017]. Chemical modifications such as N1-methylpseudouridine (m^1^Ψ) further mitigate innate immune sensing and enhance protein output [Karikó et al., 2005, Svitkin et al., 2017, Nance and Meier, 2021].

Despite the importance of each region, recent evidence strongly supports the necessity of joint optimization across all three regions. Medjmedj et al. [2025] showed that 5′-3′ UTR base-pairing interactions can reduce protein output by 2-fold despite high ribosome loading, while 5′-3′ combinations that perform well in isolation are incompatible when paired, inhibiting translation in mice [Ye et al., 2023]. Notably, identical 5′-3′ UTR combinations produced dramatically different outcomes depending on the CDS: nanoluciferase CDS yielded benchmark-level protein expression, whereas firefly luciferase CDS resulted in approximately 10-fold lower protein output, attributed to CDS-dependent changes in global mRNA secondary structure [Medjmedj et al., 2025]. In a comprehensive analysis of all 65,536 possible translation initiation sequences, Noderer et al. [2014] revealed that dinucleotide interactions at the 5^′^ UTR-CDS boundary drive up to 24.8% differences in translation efficiency, such that optimal initiation context cannot be designed independently of either UTR or CDS. These findings demonstrate that designing full-length mRNA sequences requires co-optimization, as the combinatorial design space is astronomically large even for a fixed protein, and is further magnified when incorporating UTR variability.

Recent advances in machine learning and deep learning have accelerated mRNA design optimization. LinearDesign [Zhang et al., 2023] employs dynamic programming to jointly optimize CDS structure and codon usage, achieving up to 128-fold increases in antibody titers in preclinical COVID-19 vaccine studies. For UTR design, generative adversarial networks such as UTRGAN [Barazandeh et al., 2025] can design synthetic 5^′^ UTR sequences with up to 34-fold higher predicted translation efficiency, while Smart5UTR uses a multi-task autoencoder for generative reconstruction and optimization [Tang et al., 2024]. More recently, diffusion-based frameworks have been applied to UTR generation, including mRNAutilus [Patel et al., 2025] and RNAdiffusion [Huang et al., 2024], and the 5^′^ UTR Language Model demonstrated the feasibility of transformer-based UTR generation [Chu et al., 2024]. Several recent approaches attempt full-length mRNA design. Leppek et al. [2022] developed PERSIST-seq, a high-throughput platform that systematically evaluates diverse mRNA designs varying across 5^′^ UTR, CDS, and 3^′^ UTR regions, revealing that in-cell mRNA stability is a greater driver of protein output than ribosome load alone. Deep generative models like GEMORNA [Zhang et al., 2025] employ separate transformer-based models tailored for generating 5^′^ UTR, CDS, and 3^′^ UTR sequences independently, which are then combined to create full-length optimized mRNA sequences. Similarly, integrated approaches such as iDRO [Gong et al., 2023] use BiLSTM-CRF for CDS optimization coupled with RNA-Bart for UTR generation, demonstrating full-length optimization across all three regions. For multi-objective UTR optimization, MOBO-5UTR couples a latent DNA language model with Bayesian optimization [Yamada et al., 2025], and RNop applies a deep learning framework with four specialized loss functions [Gong et al., 2025]. However, none of these approaches jointly optimize all three regions while capturing the long-range interactions across regions, which are critical to biological function, mRNA stability, and protein output.

High-throughput experiments have enabled extensive collection and curation of high-quality datasets measuring critical mRNA properties including mRNA half-life, translation rates, and protein expression across diverse sequences [Leppek et al., 2022, Sample et al., 2019, Cao et al., 2021, He et al., 2024]. For instance, Sample et al. [2019] generated a massively parallel reporter assay (MPRA) library of 280,000 random 5^′^ UTR sequences and measured mean ribosome load via polysome profiling in human cells, providing a foundational dataset for training deep learning models to predict translation efficiency from sequence. These high-quality datasets provide the foundation for building powerful predictive models and enabling further design exploration. Some methods utilize powerful predictive models trained on high-throughput sequencing (HTS) data to score and rank carefully collected sequences, which are designed by multiple algorithms and strategies. However, these approaches critically depend on the quality of the candidate pool.

To address these limitations, we propose mRNA-GPT, a generative model for *de novo* full-length mRNA sequence design and optimization. The model was pre-trained on 30 million full-length mRNA sequences with the three regions annotated to capture interactions across regions (5^′^ UTR, CDS, and 3^′^ UTR) often missed by modular approaches. We employ reinforcement learning techniques that directly and iteratively optimize the mRNA-GPT model for enhanced target properties, with flexibility to incorporate any task-specific reward model. Furthermore, we extend our framework to multi-objective optimization, simultaneously maximizing half-life and translation rate. In particular, mRNA-GPT is highly flexible during inference, enabling three different generation modes: (1) *de novo* generation of any single region; (2) full-sequence generation; or (3) conditional generation of one or two regions given any other region. Such capabilities are powered by random region ordering during pre-training.

## Methods

mRNA-GPT is a decoder-only architecture for full-length mRNA generation. The pre-training dataset consists of 30 million natural mRNA sequences collected from the NCBI Reference Sequence database (RefSeq) [Li et al., 2025, 2024]. Each sequence contains three regions: 5^′^ UTR, CDS, and 3^′^ UTR. As illustrated in Fig. 1a, the 5^′^ UTR and 3^′^ UTR are tokenized into segments using pre-trained tokenizers, while coding regions are split into sequential codons. Special tokens [5UTR], [CDS], and [3UTR] are added at the beginning of each region to indicate different functional areas, and the special token [EOS] is added after the 3^′^ UTR.

**Figure 1.**
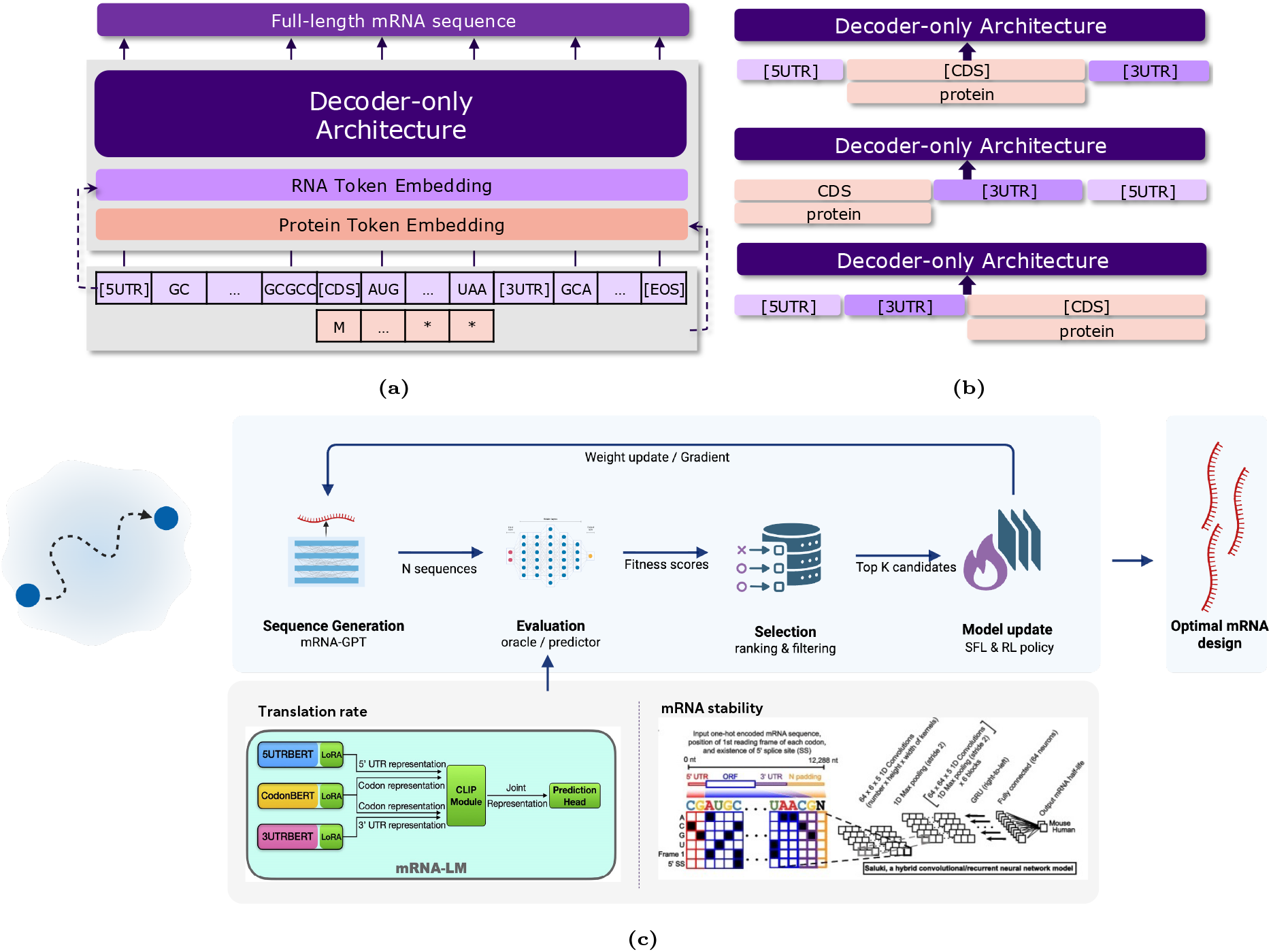
(a) Model architecture, (b) pre-training data order shuffling, and (c) iterative optimization framework. (a) mRNA-GPT is a decoder-only model for full-length mRNA design using two embedding layers: RNA and protein, and pre-trained tokenizers for UTR regions. (b) Pre-training permutes 5^′^ UTR, CDS, and 3^′^ UTR order. (c) The iterative optimization loop: the model **generates** candidate mRNA sequences, a reward model **scores** each sequence for the target property (e.g., stability or translation efficiency), the top-performing candidates are **selected**, and the generator is **updated** via supervised fine-tuning (SFT) or reinforcement learning (PPO). This cycle repeats for multiple iterations to progressively improve sequence quality.

Along the coding region, target protein sequences are incorporated to guide sequence design, ensuring that generated coding sequences can be translated into the target protein. The protein sequences consist of amino acids that are shifted left by one position to align the next amino acid with the generation of the corresponding codon. Specifically, the methionine (Met) amino acid, which corresponds to the start codon (AUG), is aligned with the special token [CDS] and fed into the model, thereby training the model to generate the start codon AUG next. Consequently, the architecture employs two input embedding layers: for coding regions, codon embeddings and amino acid embeddings are concatenated and fed into the model, while for UTR regions, only special tokens [5UTR] and [3UTR] serve as regional indicators.

Unlike simple nucleotide-level modeling or fixed n-gram tokenization for UTR regions, we trained data-driven tokenizers separately for 5^′^ UTR and 3^′^ UTR. Each tokenizer was pre-trained on the same corpus used for mRNA-GPT (30 million sequences) with a fixed vocabulary size of 5000 subword units using the SentencePiece framework [Kudo and Richardson, 2018]. Tokenization employed a subword model based on byte-pair encoding (BPE). On average, the 5^′^ UTR is approximately 200 *nt* and the 3^′^ UTR is approximately 800 *nt* in length. The use of pre-trained subword tokenizers significantly reduces the number of tokens representing these sequences, shrinking hundreds of nucleotide tokens to fewer than a hundred subword tokens. This compression enables models to process longer RNA spans within fixed context window limits, thus benefiting performance on full-length mRNA tasks. Compared to fixed-length n-gram tokenizers, pre-trained tokenizers learn variable-length tokens according to sequence frequency; these tokens can better capture meaningful regulatory motifs, whereas rigid n-gram tokenization may break motifs and result in less biologically relevant segmentation.

As shown in Fig. 1b, during pre-training, we randomly shuffle the order of the three regions to enhance scalability and enable diverse downstream tasks. Three possible orders are employed: 5^′^ UTR-CDS-3^′^ UTR, CDS-3^′^ UTR-5^′^ UTR, and 5^′^ UTR-3^′^ UTR-CDS. Such a flexible training strategy enables conditional generation capabilities, allowing the model to generate any region conditioned on other regions. For instance, beyond following the natural 5′ to 3′ order, the model can generate 5^′^ UTR given a 3^′^ UTR, CDS given both UTR regions, or UTR regions based on CDS. Besides the customized two input layers, pre-trained tokenizers, and shuffled order of regions, the core architecture utilizes a GPT-2 model [Radford et al., 2019] with a maximum length of 1024 tokens, 16 layers, 16 attention heads, and 768-dimensional token embeddings.

During inference, mRNA-GPT supports three generation modes: (1) generating a single region by prompting with the appropriate special token ([5UTR], [CDS], or [3UTR]); (2) full-length mRNA generation, in which the model sequentially generates the 5^′^ UTR, CDS, and 3^′^ UTR from 5′ to 3′, with each region conditioned on all previously generated regions; and (3) conditional generation using alternative region orderings enabled by the shuffled pre-training strategy. For full-length generation, the process proceeds in three stages: first, the 5^′^ UTR is generated token by token from the [5UTR] prompt; second, the [CDS] token and target protein sequence are appended, and the CDS is generated conditioned on the 5^′^ UTR context; finally, the [3UTR] token is appended, and the 3^′^ UTR is generated conditioned on both upstream regions, until the end-of-sequence token ([EOS]) is produced.

To optimize the model for target properties of interest, such as mRNA half-life, translation rate, or protein output, we applied an iterative optimization process. Figure 1c illustrates two alternative approaches: supervised fine-tuning (SFT) or reinforcement learning (RL) via Proximal Policy Optimization (PPO). Both approaches incorporate a reward model (RM), also referred to as the oracle, which is a supervised model trained on experimental datasets to predict a target property value for a given mRNA sequence. In this work, we employ two reward models as shown in Fig. 1c: (1) Saluki [Agarwal and Kelley, 2022], a hybrid convolutional and recurrent deep neural network trained on experimentally measured human mRNA half-lives to predict mRNA stability directly from nucleotide sequence; and (2) mRNA-LM [Li et al., 2025], a mRNA language model fine-tuned on experimentally measured human transcripts to predict translation efficiency. The reward model remains static within each optimization iteration; however, once experimental results from mRNA-GPT designs are obtained, the reward model can be updated to increase prediction accuracy as part of a larger active learning framework [Bailey et al., 2024].

The key differences between these two approaches lie in their data collection methods for fine-tuning and the optimization objectives. In the SFT approach, sequences are scored and ranked by the reward model, with only the top-performing sequences retained. The current model is then fine-tuned on these top sequences using next-token prediction loss. For a sequence of length *N* with tokens *x*_1_, *x*_2_, …, *x*_*N*_, where the model predicts the probability *p*_*θ*_(*x*_*i*_|*x*_*<i*_) of token *x*_*i*_ given its left context, averaged over sequence length (per-token loss):

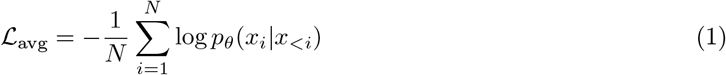

where *θ* represents the set of all trainable model parameters. Through this process, the model learns the statistical patterns of high-performing sequences and subsequently generates optimized sequences with improved property values.

The PPO strategy utilizes all sequences generated by the current model, known as the policy model. The policy model (*π*_*θ*_) refers to the currently updated model whose parameters are being optimized, while the reference model (*π*_ref_) is typically the supervised pre-trained or fine-tuned (SFT) model and is kept frozen to constrain policy updates and maintain output quality. PPO updates the policy model conservatively: smaller parameter changes per update empirically lead to better convergence and stability. To prevent excessively large policy shift, PPO computes a ratio between the current policy and the old policy (i.e., the policy at the start of the current update step):

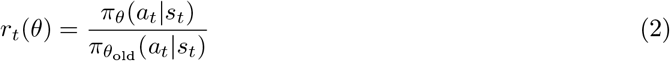

where *π*_*θ*_(*a*_*t*_|*s*_*t*_) and 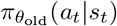 are the probabilities of taking action *a*_*t*_ in state *s*_*t*_ under the current and old policies, respectively. If *r*_*t*_(*θ*) *>* 1, the action *a*_*t*_ is more likely under the current policy than under the old policy, quantifying the magnitude of the policy shift. PPO uses the following clipped surrogate objective to ensure a conservative update:

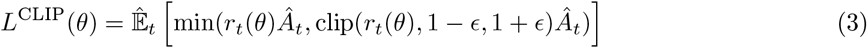

where *Â*_*t*_ is the advantage function, representing how much better an action is compared to the expected value for a given state. In practice, advantage values are often computed from normalized rewards with a baseline value. The “min” operator provides a pessimistic (lower bound) estimate: it takes either the clipped or unclipped objective depending on whether the update would exceed the allowed *ϵ*-range (typically *ϵ* = 0.2). Additionally, a Kullback-Leibler (KL) divergence penalty between the current policy and the frozen reference model is computed:

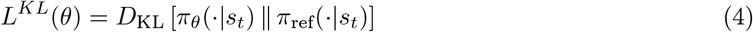

The complete loss is

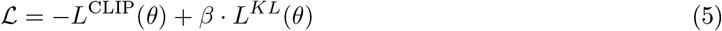

where *β* is the KL penalty coefficient. Training maximizes the clipped surrogate objective while minimizing the KL divergence, thus balancing reward maximization with a constraint on divergence from the reference policy, which is kept frozen during optimization.

## Results

### De Novo Generation of Diverse 3’ UTRs with Enhanced Stability

The 3^′^ UTR is a principal determinant of mRNA stability: it harbors AU-rich elements, microRNA binding sites, and RNA-binding-protein recognition motifs that collectively govern transcript half-life in the cytoplasm [Mayr, 2017, 2019]. Because stability is often the rate-limiting factor for protein output [Leppek et al., 2022], optimizing the 3^′^ UTR is a high-priority objective in mRNA therapeutic design. Accordingly, we use the Saluki model [Agarwal and Kelley, 2022] as the reward model (see Methods) to predict mRNA half-life and guide iterative optimization of 3^′^ UTR sequences. Among the most relevant baselines, GEMORNA [Zhang et al., 2025] introduced GEMORNA-UTR, a decoder-only transformer architecture pre-trained on natural UTR sequences (both 5^′^ UTR and 3^′^ UTR). For 3^′^ UTR design, GEMORNA-UTR was fine-tuned on top-performing natural sequences scored by PRED-3UTR, a predictive model trained on 90 000 experimentally labeled 3^′^ UTR sequences with measured degradation rates. This approach is limited by reliance on natural sequences and a single fine-tuning step, increasing the risk of local optima and restricting sequence diversity. In contrast, mRNA-GPT utilizes an iterative optimization strategy with an in-the-loop reward model (see Methods), enabling direct optimization of desired properties. Our model eliminates the need for seed or natural sequences, evolving the candidate pool entirely through self-generated designs.

Both the publicly available GEMORNA model and the mRNA-GPT model (prompted with the [3UTR] token) were evaluated across predefined length bins. For the GEMORNA model, 10 000 sequences were generated per bin; after removing duplicates, 128, 119, and 121 unique 3^′^ UTR sequences remained in the short (35-65 *nt*), medium (66-95 *nt*), and long (96-125 *nt*) length bins, respectively. Note that the released GEMORNA model produces only about 100 diverse sequences per bin; therefore, its samples may not be representative. For both the pre-trained mRNA-GPT and fine-tuned mRNA-GPT, 5000 unique 3^′^ UTR sequences were generated per length bin. Because mRNA-GPT can produce arbitrarily long sequences, an additional extra-long bin (126-200 *nt*) with 5000 sequences was included. As a natural baseline, 1000 vertebrate 3^′^ UTR sequences from RefSeq were collected per bin, yielding 4000 reference sequences.

Comparing sequence properties across samples, the length distribution of natural 3^′^ UTR sequences is broad and relatively uniform (Fig. S2a), whereas both GEMORNA and fine-tuned mRNA-GPT samples exhibit greater variability, with distinct peaks suggesting preferential generation within certain length ranges. In terms of nucleotide composition (Fig. 2d), natural and GEMORNA sequences have uniform nucleotide distributions, whereas pre-trained and fine-tuned mRNA-GPT samples are cytosine-rich. Notably, across optimization iterations, cytosine content increases while guanine content decreases. This shift is consistent with two known stabilizing mechanisms in 3^′^ UTR regions: (1) C-rich sequences promote binding of poly(C)-binding protein 2 (PCBP2) and *α*-complex formation, which shields transcripts from exonucleolytic decay [Ji et al., 2013]; and (2) reduced guanine content lowers the propensity for G-quadruplex formation, secondary structures in the 3^′^ UTR that can recruit destabilizing factors and accelerate mRNA degradation [Lee et al., 2020]. Fine-tuning lowers the overall GC content compared to pre-trained mRNA-GPT, which is consistent with the known negative correlation between GC content and stability [Litterman et al., 2019]. Length-normalized minimum free energy (MFE), calculated with the ViennaRNA package [Lorenz et al., 2011] (see Eq. (S1)), shows that fine-tuned mRNA-GPT sequences have lower MFE values, indicating greater thermodynamic stability and closer resemblance to natural profiles (Fig. 2e) [Lorenz et al., 2011, Zhang et al., 2023]. Similarly, melting temperature (Tm) analysis (Fig. 2f; see Eq. (S2)) reveals that fine-tuning increases thermal stability to levels comparable to natural and GEMORNA sequences [Cock et al., 2009].

**Figure 2.**
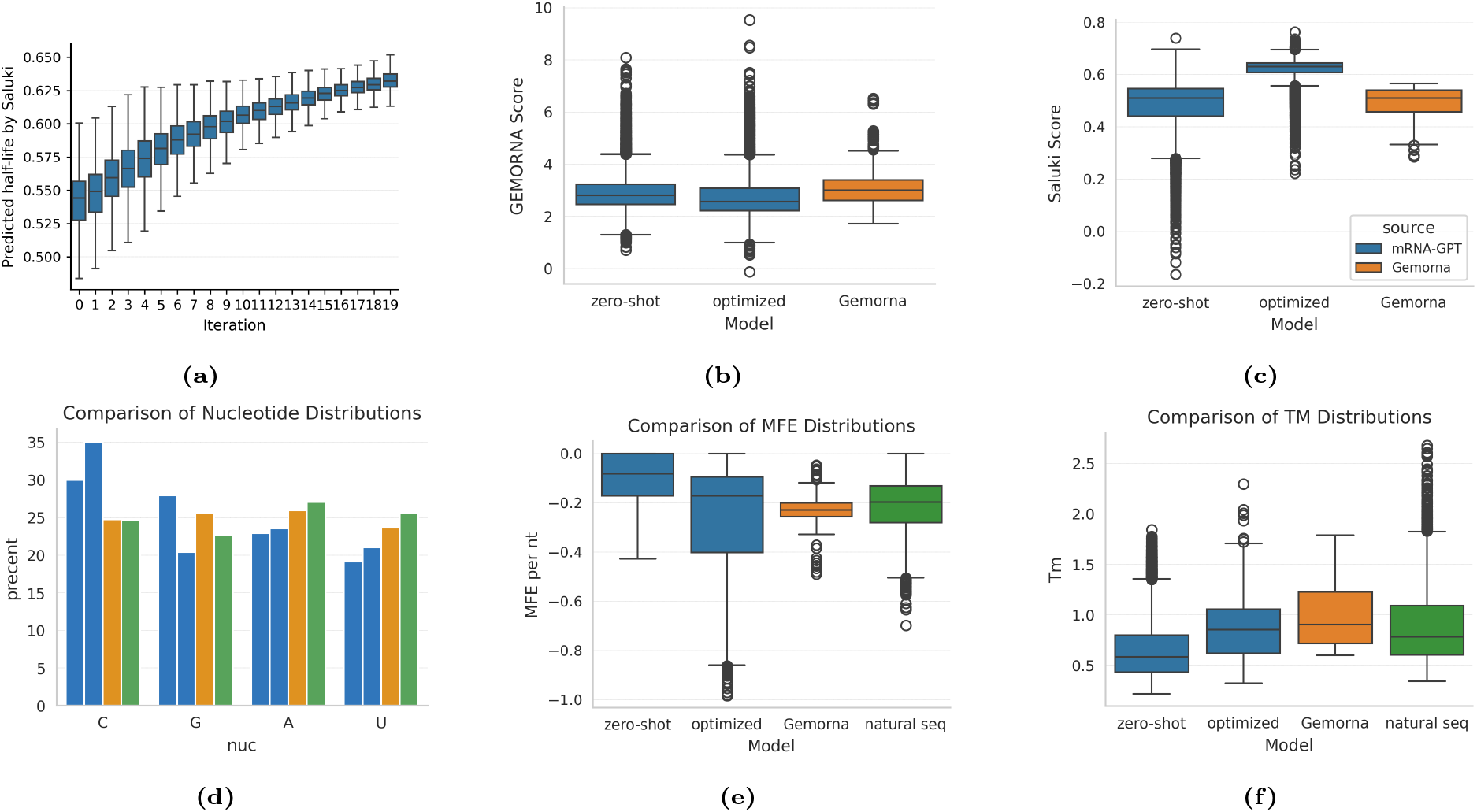
3^′^ UTR optimization, analysis, and benchmarking. (a) Box plot of optimization progression showing predicted half-life improvements across iterations for the short-length bin. (b) Benchmarking using the GEMORNA PRED-3UTR oracle. (c) Benchmarking using the Saluki oracle. Bar plots of average nucleotide composition. (e) Violin plots of MFE. (f) Box plots of melting temperature.

To assess novelty, we converted the pre-trained dataset of over 30 million 3^′^ UTR sequences into a BLAST database using a 15-*nt* threshold for hits and measured each sequence’s longest match (Fig. S2b). Fine-tuned mRNA-GPT produces a greater number of novel sequences than the pre-trained mRNA-GPT. To assess diversity, we computed 5-mer frequencies for each sequence in the fine-tuned mRNA-GPT and then calculated cosine similarity (see Eq. (S3)). We find that the mean cosine similarity is 0.23, indicating high diversity and broad sequence space exploration with minimal overlap in k-mer patterns. Collectively, these results suggest improved predicted stability in the fine-tuned mRNA-GPT outputs. The fine-tuned mRNA-GPT performs on par with GEMORNA and natural sequences, occasionally surpassing them. It also produces a higher proportion of novel and diverse 3^′^ UTR sequences, demonstrating its capacity to generate diverse, novel sequences with favorable biophysical properties.

We applied iterative optimization via PPO for 20 iterations, sampling 5000 candidate 3^′^ UTR sequences per bin each round. Fig. 2a demonstrates improved predicted half-lives across iterations (shown for the short bin), evidencing the effectiveness of the iterative optimization procedure. Notably, the median predicted half-life increased from 0.54 to 0.63, and the maximum value from 0.63 to 0.67. Corresponding results for all four bins are in Fig. S1. Fig. 2c displays the predicted half-life distributions for sequences generated by mRNA-GPT, GEMORNA-UTR, and natural samples. The pre-trained mRNA-GPT achieves a comparable median to GEMORNA-UTR and natural sequences, but spans a wider range, especially for longer sequences, thus producing some sequences with higher predictions than GEMORNA-UTR. Fine-tuning mRNA-GPT further improves predicted half-lives beyond all benchmarks. For a direct comparison, sequences from each method were also scored using the PRED-3UTR oracle used in GEMORNA-UTR optimization (Fig. 2b). In this setting, pre-trained mRNA-GPT has a lower median than GEMORNA-UTR but exhibits many high-value outliers, outperforming GEMORNA-UTR at the distribution tail. Fine-tuned mRNA-GPT is not directly optimized to maximize predictions by the PRED-3UTR oracle and therefore does not achieve high predicted values under this metric, which reflects the specificity of the optimization target. These results demonstrate that mRNA-GPT’s iterative, reward-driven optimization enables broader exploration and superior identification of high-stability 3^′^ UTR designs.

### Oracle-Guided CDS Optimization Achieves Superior Translation Rates While Balancing Codon Adaptation, Structural Stability, and Diversity

We next applied the iterative SFT optimization framework (see Methods) to coding sequence design, training a conditional generator *p*(CDS |protein) to produce CDS with progressively higher predicted translation rates, as scored by the reward model mRNA-LM [Li et al., 2025]. We optimized the CDS regions of mRNA sequences corresponding to four target proteins, as listed in Table 1. At each iteration *t* (with *t*=0 representing the pre-trained model), the generator produces 6000 candidate CDS sequences per protein. All candidates were scored by the reward model, generating a ranked list per protein and iteration. The top *N* =1500 highest-scoring CDS sequences from each list were used as the fine-tuning dataset for that protein at the given iteration. For *t* ≥1, sampling and fine-tuning were repeated from the previous iteration’s checkpoint, enabling each round to refine a protein-specific model using the best sequences identified thus far.

**Table 1:** Target proteins used in CDS design and optimization.

| ID | Organism | Protein name | Accession | Length (aa) | Brief description |
| --- | --- | --- | --- | --- | --- |
| 1 | Rabies virus (strain Pasteur vaccins / PV) (RABV) | Glycoprotein | P08667 | 524 | Trimeric viral fusion glycoprotein in the rabies virion envelope; main target of neutralizing antibodies. |
| 2 | Zaire ebolavirus (strain Eckron-76) (ZEBOV) (Zaire Ebola virus) | Envelope glycoprotein | P87671 | 676 | Heavily glycosylated class I envelope glycoprotein cleaved into GP<sub>1</sub>/GP<sub>2</sub>; mediates host-cell attachment and membrane fusion. |
| 3 | Severe acute respiratory syndrome coronavirus 2 (2019-nCoV) (SARS-CoV-2) | Membrane protein | P0DTC5 | 222 | Multi-pass structural protein that shapes the coronavirus envelope and scaffolds virion assembly. |
| 4 | Homo sapiens (Human) | L-dopachrome tautomerase | P40126 | 519 | Melanosomal enzyme catalyzing the conversion of L-dopachrome to DHICA in the melanogenesis pathway. |

We focus on a representative target, the SARS-CoV-2 membrane protein (Protein 3 in Table 1), with results for the remaining three proteins listed in Table 1 (amino acid sequences in Supplementary Table S1) provided in the Supplementary Information. For benchmarking, we included both a classical algorithm (LinearDesign) and the state-of-the-art language model-based GEMORNA as baselines. GEMORNA-CDS, an encoder-decoder module specifically designed for CDS design, treats protein sequences as the source and mRNA CDS as the target. We used GEMORNA-CDS to generate 200 unique CDS sequences for each protein. For LinearDesign, we systematically evaluated a range of *λ* values (from 0 to 100 in steps of 0.1) to balance CAI and MFE. For the SARS-CoV-2 membrane protein, setting *λ* = 10 produced a CDS with a CAI value of 0.991. After deduplication, this yielded 37 unique CDS sequences.

Fig. 3a presents the distribution of predicted translation rates for the eleven iterations, alongside the LinearDesign and GEMORNA-CDS baselines. The pre-trained model (*t*=0) produces a broad score distribution with a median near 1.0 and a pronounced left tail containing sequences with negative predicted translation rates. As optimization proceeds, the distribution shifts upward: medians increase nearly monotonically, upper quartiles rise, and the interquartile range narrows. By iteration *t*=10, low-scoring outliers are almost eliminated and the median score has approximately doubled compared to *t*=0. The final iterations surpass both baselines. The GEMORNA-CDS distribution lies below mRNA-GPT outputs from rounds *t*=7–10 in both median and upper quantiles. LinearDesign performs better than GEMORNA-CDS but remains slightly below mRNA-GPT at rounds *t*=9–10, where mRNA-GPT achieves a higher median and 75th percentile. Note that the GEMORNA-CDS model uses zero-shot generation; thus its sequences provide a strong starting point but lack further optimization. In contrast, mRNA-GPT’s iterative oracle-guided optimization yields continued performance improvements, surpassing both baselines and demonstrating superior capacity for translation rate enhancement.

**Figure 3:**
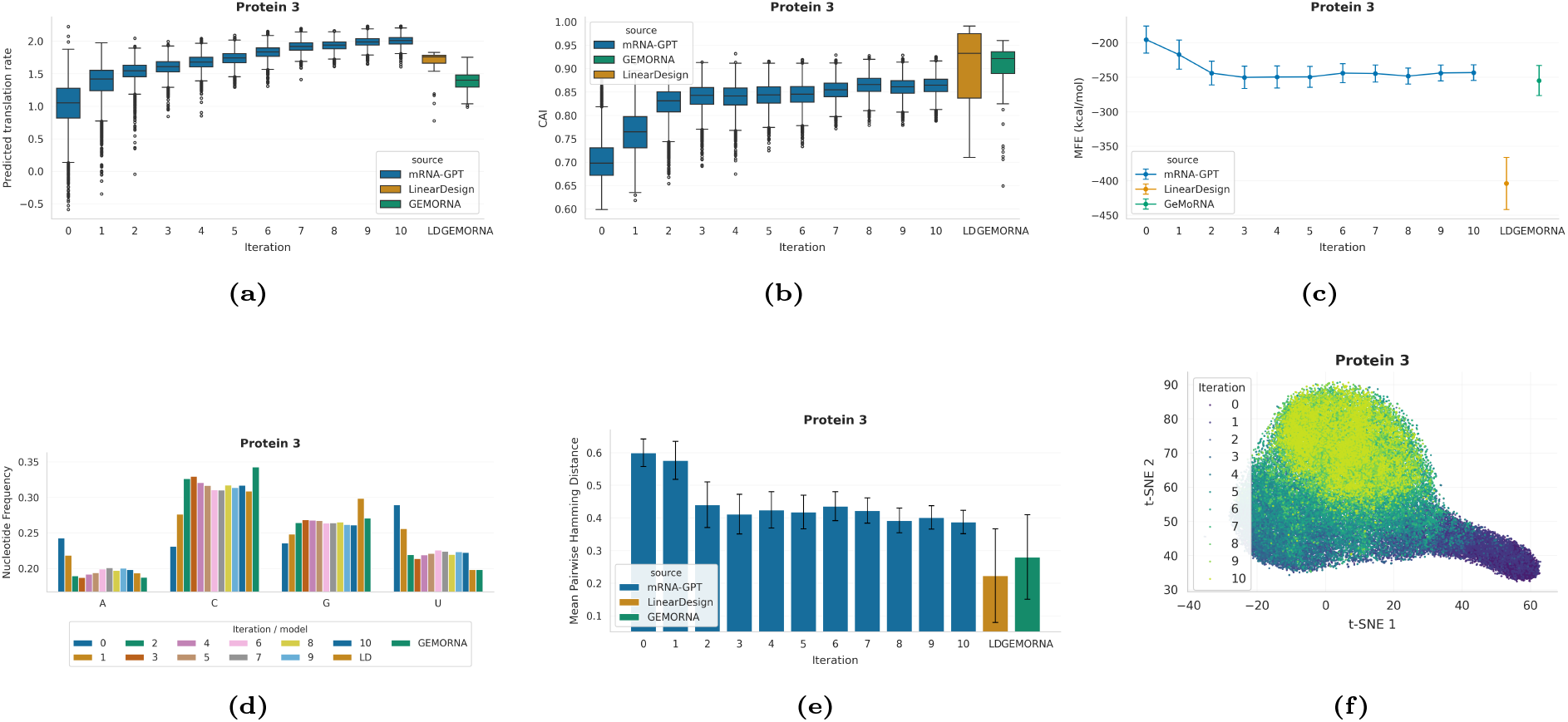
CDS optimization readouts for SARS-CoV-2 membrane protein. Panels (a)–(f) depict properties of synonymous CDS generated by mRNA-GPT across iterations and by the LinearDesign and GEMORNA baselines: (a) predicted translation rates, (b) CAI, (c) MFE, (d) mean nucleotide composition, (e) sequence diversity, and (f) t-SNE embedding of codon-usage vectors.

Fig. 3b reports the codon adaptation index (CAI) distributions, computed from a fixed human codonusage reference table following the classical definition of codon adaptation index [Sharp and Li, 1987] (see Eq. (S5)). At *t*=0, the pre-trained mRNA-GPT shows CAI values centered around ∼0.70, indicating modest adaptation to human codon usage with room for improvement. Over ten optimization rounds, the CAI distribution drifts upward in an almost monotonic fashion, reaching medians around ∼0.87 at *t*=10 with tighter interquartile ranges and fewer low-CAI outliers. Oracle-guided fine-tuning therefore implicitly pushes the generator toward human-preferred codons, despite CAI not being optimized explicitly. Both LinearDesign and GEMORNA achieve even higher CAI values than *t*=10 mRNA-GPT, with medians in the ~0.9–0.95 range. However, this higher CAI does not yield superior predicted translation rates (Fig. 3a), in line with the view that codon optimality contributes to, but does not solely determine, translation efficiency and mRNA stability [Presnyak et al., 2015, Wu et al., 2019]. This suggests that aggressively maximizing codon adaptation alone is not optimal for the oracle and that mRNA-GPT converges to a more balanced region of sequence space.

We next examine global RNA secondary structure using ViennaRNA `RNAfold` [Lorenz et al., 2011] to compute the Minimum Free Energy (MFE) for the designed CDS sequences. Fig. 3c plots the mean MFE (kcal/mol) and its standard deviation over generated sequences, with more negative values indicating more stable structures. For the pre-trained mRNA-GPT (*t*=0), the mean MFE is relatively shallow (around −200 kcal/mol), indicating comparatively unstable transcripts. Early optimization steps drive MFE downwards: by *t*≈2 the mean MFE has decreased to roughly −240 to −250 kcal/mol. Beyond *t*≈3, the curve plateaus with only small fluctuations. Optimization therefore follows a “stabilize-then-saturate” trajectory: transcripts become substantially more stable than at *t*=0, but later iterations preserve this stability rather than pushing toward extreme over-folding. LinearDesign produces much more stable (more negative) structures, with mean MFE markedly lower than any mRNA-GPT iteration, reflecting its explicit energy-minimization objective [Zhang et al., 2023]. GEMORNA lies closer to the *t*=10 mRNA-GPT regime, with similar or slightly more negative MFEs. Combined with the translation-rate results, this indicates a compromise between stability and accessibility: mRNA-GPT transcripts are more structured than the pre-trained baseline yet less over-stabilized than energy-minimizing designs, while achieving higher predicted translation.

Fig. 3d complements this view by summarizing nucleotide frequencies (T→U). The pre-trained model starts with relatively modest GC content, with A and U over-represented compared to C and G. Across iterations, optimization progressively enriches C and G while depleting A and U, so that by *t*=10 the GC fraction closely matches that of both LinearDesign and GEMORNA. The model thus learns to increase GC content, a known correlate of mRNA stability and expression in multiple systems [Presnyak et al., 2015, Zhang et al., 2023, 2025], but only to the extent supported by the oracle, without overshooting the regime explored by codon-centric baselines.

To monitor potential mode collapse under top-*k* selection, we quantify diversity as the mean pairwise Hamming distance (the fraction of codon positions at which two sequences differ), normalized by CDS length, across generated samples based on a random sample of sequence pairs per group (Fig. 3e). The pre-trained model exhibits high diversity, with mean pairwise distances around ∼0.6. Optimization compresses the distribution: by *t*≈2–3 the mean distance has decreased to ∼0.4, then fluctuates mildly around this value. Even after ten rounds, mRNA-GPT maintains substantially higher diversity than both LinearDesign and GEMORNA, whose mean distances cluster around ∼0.2–0.3. Thus, while low-scoring regions of sequence space are pruned, the model continues to explore a wide variety of synonymous CDS solutions rather than collapsing onto a single codon pattern, an important requirement for robust mRNA design.

A t-distributed stochastic neighbor embedding (t-SNE) of codon-usage vectors (Fig. 3f) provides a complementary view [van der Maaten and Hinton, 2008]. Each sequence is represented by a 64-dimensional, L1-normalized codon-usage vector, and t-SNE is applied to sets of these vectors. Points are colored by optimization iteration. The embedding reveals a broad, continuous manifold: early iterations (*t*=0–1) populate one region, while later iterations gradually drift toward other areas along a smooth trajectory. Late iterations still span a wide part of the manifold, consistent with the diversity analysis and indicating that optimization reweights, rather than collapses, codon-usage modes.

Analogous analyses for Rabies virus G, Ebola GP, and human DCT/TYRP2 are provided in the Supplementary Material. Qualitatively, they show the same hallmarks as SARS-CoV-2 membrane protein: batch-wide shifts toward higher oracle scores, improved CAI and GC content, moderate compaction but no catastrophic loss of diversity, and a balance between increased structural stability and avoidance of extreme over-folding (Fig. S3 to S8). Overall, our iterative mRNA-GPT optimization procedure appears to converge toward a human-expression-optimized sequence manifold that outperforms codon-centric baselines while retaining a rich set of synonymous CDS solutions, in line with the goals of recent deep generative frameworks for mRNA optimization [Zhang et al., 2025, Khoroshkin et al., 2024, Riley et al., 2025, Gong et al., 2023, Patel et al., 2025].

### End-to-End Full-Length mRNA Design Outperforms Regional Optimization

Building upon our evaluation of mRNA-GPT’s regional optimization capabilities, we next assessed the model’s ability to generate complete full-length mRNA sequences. We employed the stepwise autoregressive generation procedure described in Methods and illustrated in Fig. 4a, designing mRNA sequences from the 5′ to 3′ end with each region conditioned on all previously generated regions.

**Figure 4:**
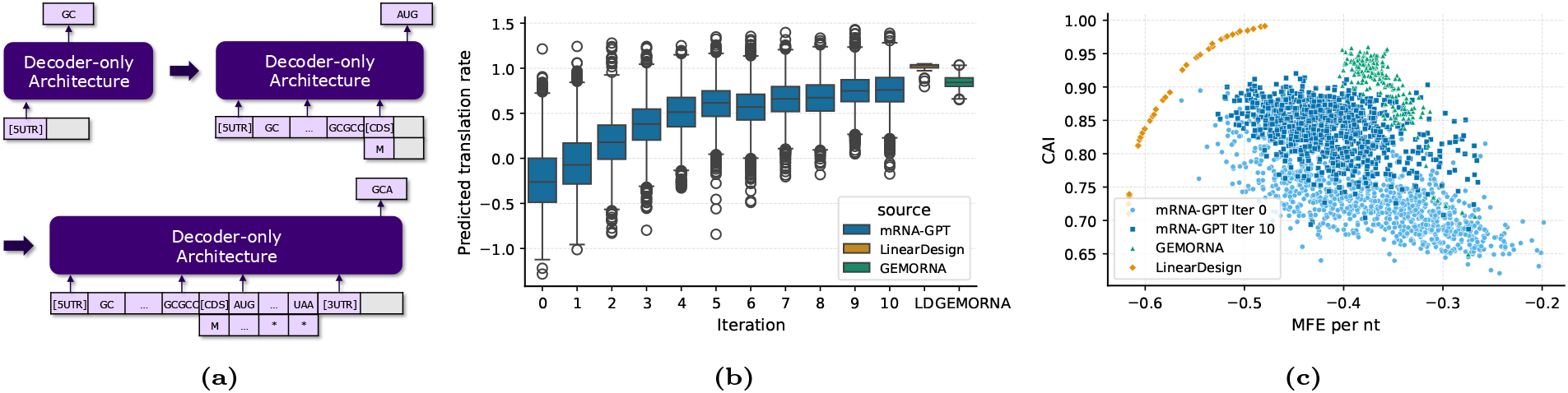
End-to-end Optimization of Full-length mRNA sequences. (a) Stepwise autore-gressive generation of 5^′^ UTR, CDS, and 3^′^ UTR. (b) Iterative optimization with mRNA-GPT yields enhanced translation rates, alongside GEMORNA and LinearDesign baselines. (c) CAI versus MFE for sequences generated by different methods.

To optimize full-length mRNA sequences, we employed mRNA-LM as the reward model to predict translation efficiency given a complete mRNA sequence [Li et al., 2025]. We applied iterative supervised fine-tuning using mRNA-GPT to optimize sequences encoding the SARS-CoV-2 membrane protein with 222 amino acids. Fig. 4b demonstrates progressive improvement in predicted translation efficiency over 10 optimization iterations. In each iteration, mRNA-GPT generated 6,000 sequences. The average predicted translation efficiency increased from -0.24 to 0.76, with convergence observed after approximately five iterations. Mann-Whitney–Wilcoxon tests [Mann and Whitney, 1947] confirmed that each iteration (except iteration 6) yielded sequences with statistically significantly higher translation efficiency than the previous iteration (p<0.05).

We benchmarked our results against two established approaches: GEMORNA and LinearDesign. For GEMORNA, we generated 5′ and 3^′^ UTR sequences independently using GEMORNA-UTR, scoring each with mRNA-LM and retaining the top 10 sequences per UTR region. We additionally generated 200 unique CDS sequences using GEMORNA-CDS. The combination of these regions yielded 10×10×200 = 20, 000 full-length sequences. The predicted translation rate distribution for GEMORNA ranged from 0.65 to 1.04 with an average value of 0.85. Mann-Whitney–Wilcoxon tests revealed that the top 10% sequences from mRNA-GPT iteration 7 to 10 exhibited significantly higher translation efficiency than the top 10% of GEMORNA sequences. As noted for isolated CDS design, GEMORNA relies on zero-shot generation without further optimization and builds separate models for UTR and CDS, thereby neglecting interactions across regions. For LinearDesign, the 37 unique CDS sequences were combined with UTR sequences from the original LinearDesign study. The resulting predicted translation rate distribution ranged from 0.79 to 1.05 with an average value of 1.01. In the final iteration (10th), the top 10% of mRNA-GPT sequences demonstrated significantly higher translation efficiency compared to the top 10% of LinearDesign sequences.

We further computed CAI (for the CDS region only) and MFE (for the full-length sequences) and visualized their relationship in Fig. 4c. Because mRNA-GPT and GEMORNA generate UTR sequences with variable lengths, we normalized the MFE by the number of nucleotides in the sequence. The figure includes only samples from the pre-trained and the last-iteration mRNA-GPT models and shows a strong negative correlation between CAI and MFE per *nt* (Pearson *r* = −0.7). The negative correlation agrees with the fundamental trade-off between MFE and CAI [Zhang et al., 2023]. Sequences designed by LinearDesign delineate the boundary of the search space because the algorithm directly optimizes both objectives simultaneously. The sequences from the optimized mRNA-GPT shift towards the top left corner with higher CAI and lower MFE compared to the pre-trained mRNA-GPT model, indicating that optimization proceeds in the intended direction. Fig. S9a and Fig. S9b show the increased CAI and decreased MFE over iterations. The broader distribution of CAI and MFE, as shown in Fig. 4c, suggests that mRNA-GPT explores a large region of the space beneath the boundary curve traced by LinearDesign; for instance, the model exploits alternative codon choices that may confer translation advantages beyond simple codon frequency optimization. In contrast, GEMORNA sequences explore a relatively small region in the upper-right corner, with higher CAI but also higher (less negative) MFE compared to mRNA-GPT, indicating that the GEMORNA model implicitly favors CAI over MFE and does not capture interactions across regions.

Interestingly, we observed contrasting optimization trajectories when optimizing 5^′^ UTR sequences in isolation versus jointly optimizing full-length mRNA sequences [Chappell et al., 2006]. When the oracle model evaluated 5^′^ UTR sequences alone, it preferentially selected shorter sequences, consistent with the principle that minimizing ribosome scanning distance enhances translation initiation efficiency. However, when optimizing complete mRNA sequences containing 5^′^ UTR, CDS, and 3^′^ UTR regions together, the optimized model (in the 10th iteration) generated significantly longer UTR sequences than the pre-trained model, as shown in Fig. S9c. This apparent contradiction reveals the importance of context-dependent optimization in mRNA design.

Moreover, full-length mRNA sequences, which include both 5^′^ UTR and 3^′^ UTR, consistently yield lower predicted translation rates compared to the CDS region alone, as shown in Fig. 4b and Fig. 3a. This reflects the critical influence of UTR regions and additional sequence context on overall translation rate. Numerous studies demonstrate that UTR regions impact ribosome recruitment, initiation, elongation, and transcript stability, often dampening the translation rate relative to optimally designed CDS-only constructs. Specifically, additional regulatory elements and increased sequence length can introduce secondary structures, regulatory motifs, and context-dependent interactions that modulate translation efficiency; for instance, longer or structured UTR can inhibit ribosome loading or scanning, and 3^′^ UTR may encode decay signals or binding sites for inhibitory factors. This context-dependent effect underscores the need for comprehensive design and optimization: considering full-length mRNA, rather than isolated CDS, is essential for realistic and biologically relevant predictions and performance in mRNA therapeutics and synthetic biology applications.

### Multi-Objective Optimization Improving Stability-Translation Trade-Offs

Optimizing UTR sequences often requires balancing multiple competing objectives, as enhancing one property (e.g., stability) can negatively impact another (e.g., translation efficiency) [Leppek et al., 2022]. Multi-objective optimization addresses these trade-offs by advancing the set of Pareto-optimal solutions, i.e., the best compromises across all objectives [Gunantara, 2018]. Here we show that a simple linear scalarization-based approach achieves competitive multi-objective performance relative to the recent methods discussed in the Introduction.

Multi-objective optimization for short (35-65 nt) 3^′^ UTR sequences was performed by iteratively updating the policy model using PPO. Optimization balanced two reward objectives: Saluki-predicted mRNA half-life (*r*_1_) and mRNA-LM-predicted translation efficiency (*r*_2_), as described in Methods. Each reward was normalized by its mean and standard deviation. The combined reward was defined as *r* = *α* · *r*_1_ + (1 − *α*) · *r*_2_ where *α* is a prior weight chosen from 0.0, 0.1, 0.5, 0.7, 0.9. For each *α*, we performed 15 iterations of the sample-score-update cycle, generating an ensemble of fine-tuned models that collectively approximated the Pareto front for stability and translation efficiency.

Hypervolume was assessed across fine-tuning iterations. Joint optimization yields distinct models that consistently dominate earlier Pareto fronts. As shown in Fig. 5e, Pareto front quality improved substantially from iteration 0 to iteration 14, with a 10.04% hypervolume increase, demonstrating enhanced model capacity for better trade-offs among objectives. The hypervolume indicator is defined as: HV(*P, r*) = *λ* (U_*p*∈*P*_ [*p, r*]), where *P* is the set of non-dominated solutions (Pareto front), *r* is the reference point, [*p, r*] denotes the hyperrectangle bounded by solution *p* and reference point *r*, and *λ* is the Lebesgue measure (volume).

**Figure 5:**
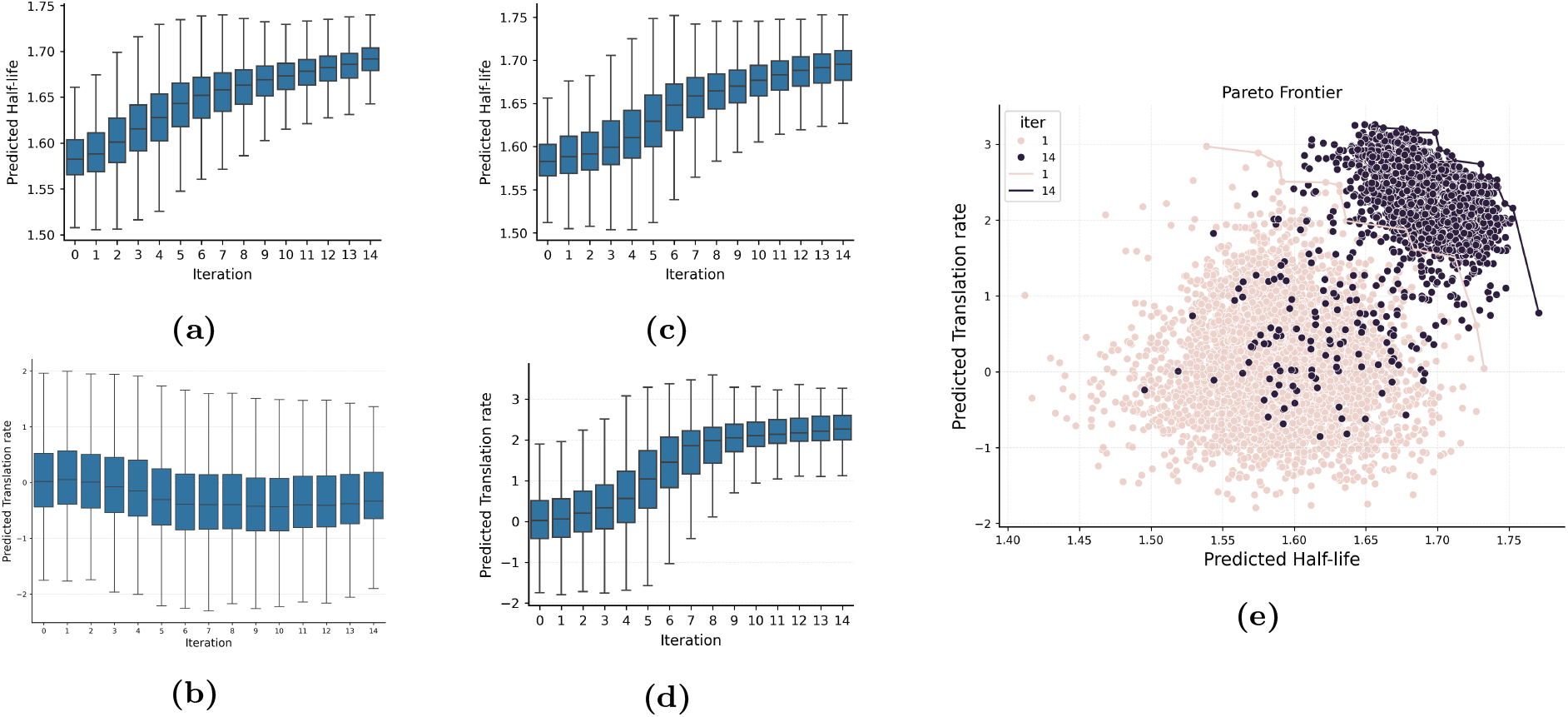
Multi-Objective Optimization. (a, c) Predicted stability (half-life) for single-objective (*α* = 0.0) and multi-objective (*α* = 0.5) optimization, respectively. (b, d) Predicted translation rate for the same strategies. (e) Pareto frontier at iterations 1 (light) and 14 (dark), showing that the optimized ensemble achieves improved trade-offs between stability and translation efficiency.

Using a joint optimization weight of *α* = 0.5 (Fig. 5c), we achieved stability improvements comparable to those from optimizing exclusively for stability (*α* = 0.0; Fig. 5a). Importantly, the joint approach resulted in a marked increase in translation efficiency (Fig. 5d), whereas optimizing solely for stability reduced translation scores (Fig. 5b). Statistical analysis with the Mann-Whitney U test indicated that multi-objective optimization (*α* = 0.5) incurred a modest decrease in stability relative to single-objective optimization (*α* = 0.0; *U* = 11,414,355, *p <* 0.0001, Cohen’s *d* = 0.15), while providing a substantial gain in translation efficiency (*U* = 137,632, *p <* 0.0001, Cohen’s *d* = 4.65). Together, these findings validate and underscore the effectiveness of our multi-objective optimization strategy.

## Conclusion

mRNA-GPT addresses a critical limitation in mRNA design: the failure to jointly optimize all functional regions while capturing long-range interactions. By pre-training on 30 million full-length sequences with randomized region ordering, our model learns complex inter-dependencies between 5^′^ UTR, CDS, and 3^′^ UTR. Iterative optimization through PPO and supervised fine-tuning enables direct maximization of target properties while maintaining sequence diversity. Empirical validation shows superior performance in 3^′^ UTR optimization, with enhanced thermodynamic stability and novelty, and in CDS optimization across four proteins, where mRNA-GPT achieves higher predicted translation rates than specialized algorithms. Full-length optimization demonstrates that optimal UTR combinations are context-dependent, confirming the necessity of joint design. The model’s flexible generation capabilities position mRNA-GPT as a versatile platform for rational mRNA design. To this end, mRNA-GPT represents a new paradigm for nucleic acid design that could accelerate development across diverse therapeutic modalities including protein replacement, antibody production, and gene editing.

## Supplementary Material

The normalized minimum free energy (MFE) is defined as:

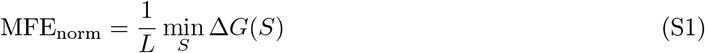

where *L* is the length of the RNA sequence (in nucleotides), *S* represents a possible secondary structure of the RNA, and Δ*G*(*S*) is the total free energy of structure *S* (in kcal/mol) as computed by ViennaRNA RNAfold.

We report length-normalized Tm values (*T*_*m*,norm_ = *T*_*m*_*/L*), where *T*_*m*_ is defined as:

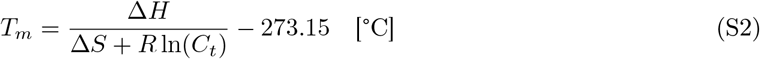

where Δ*H* and Δ*S* are the enthalpy and entropy changes from nearest-neighbor parameters, *R* is the gas constant, and *C*_*t*_ is the strand concentration.

The cosine similarity between two sequences *s*_*i*_ and *s*_*j*_ is defined as:

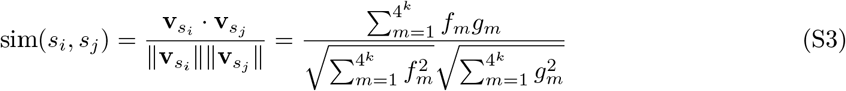

where *f*_*m*_ and *g*_*m*_ are the frequencies of the *m*-th k-mer in sequences *s*_*i*_ and *s*_*j*_, respectively. The overall diversity of a sequence set is computed as 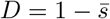, where 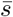 is the mean pairwise cosine similarity.

For CAI, we use codon weights

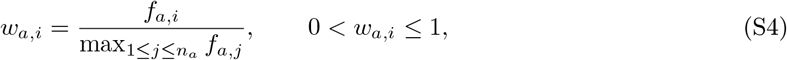

for the *i*-th synonymous codon of amino acid *a* with usage fraction *f*_*a,i*_ among its *n*_*a*_ synonymous codons. The per-sequence CAI for a CDS *g* of length *L* codons *c*_1_, …, *c*_*L*_ is then

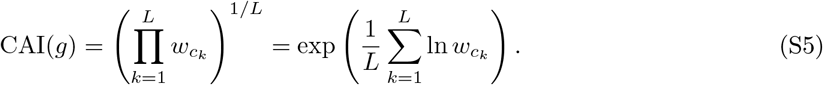

**Figure S1:**
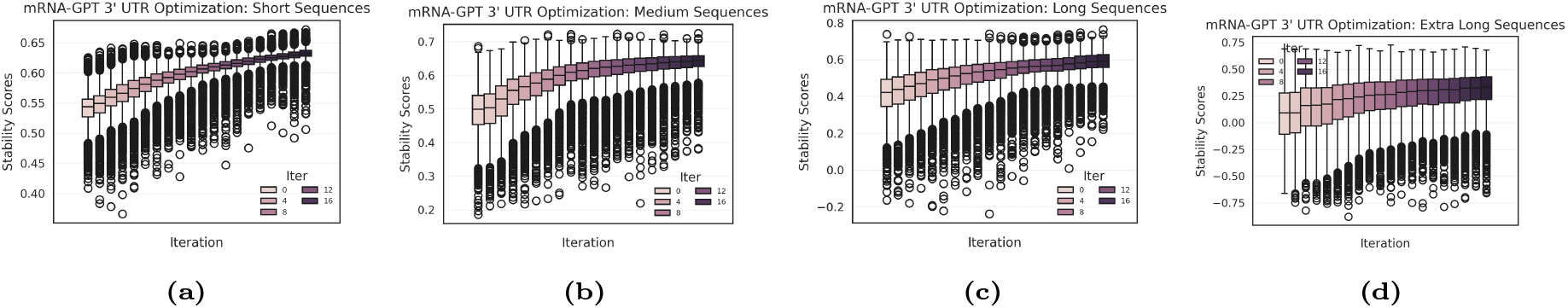
Box plot of 3^′^ UTR optimization progression and stability score improvements for short, medium, long, and extra long bin, respectively.

**Figure S2:**
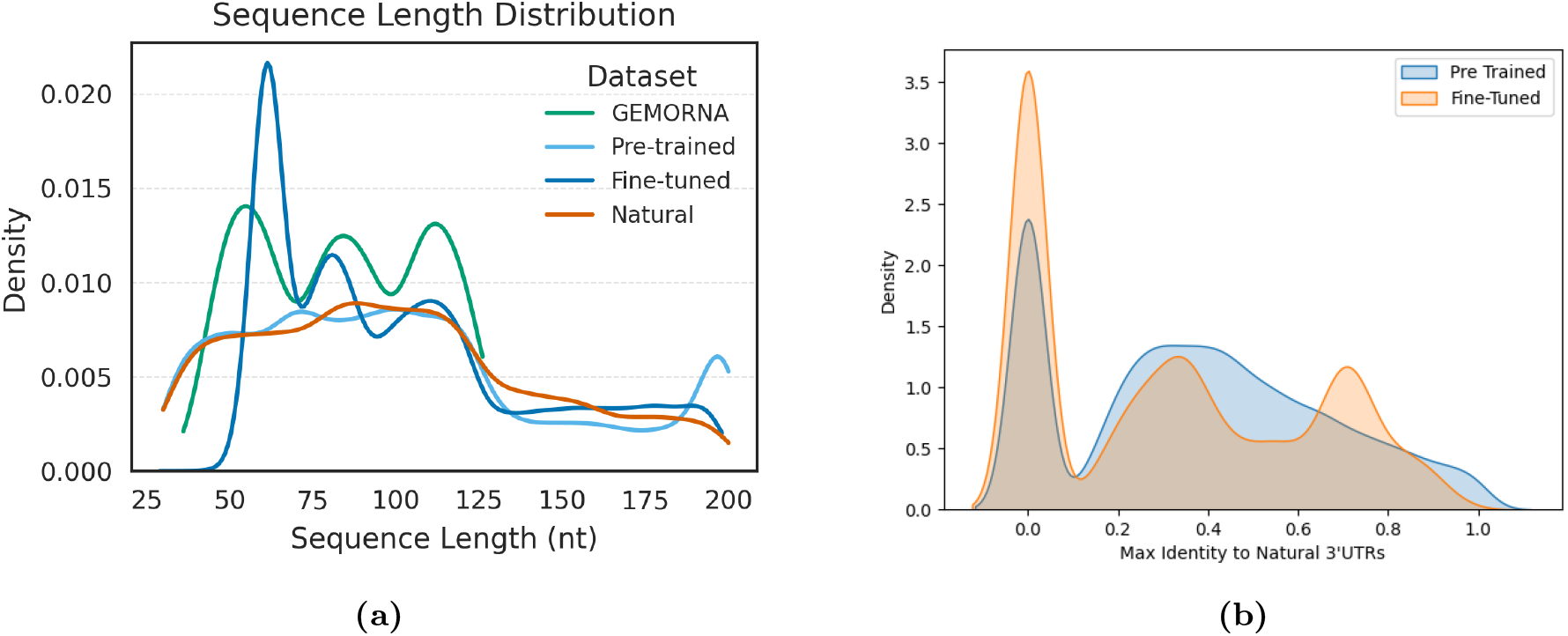
Sequence length distribution and novelty of generated 3^′^ UTR sequences. (a) Length distributions of 3^′^ UTR sequences generated by mRNA-GPT (pre-trained and fine-tuned), GEMORNA, and natural vertebrate sequences. (b) Maximum sequence identity of mRNA-GPT-generated 3^′^ UTR sequences to natural 3^′^ UTR sequences in the pre-training dataset, assessed via BLAST.

**Table S1:**
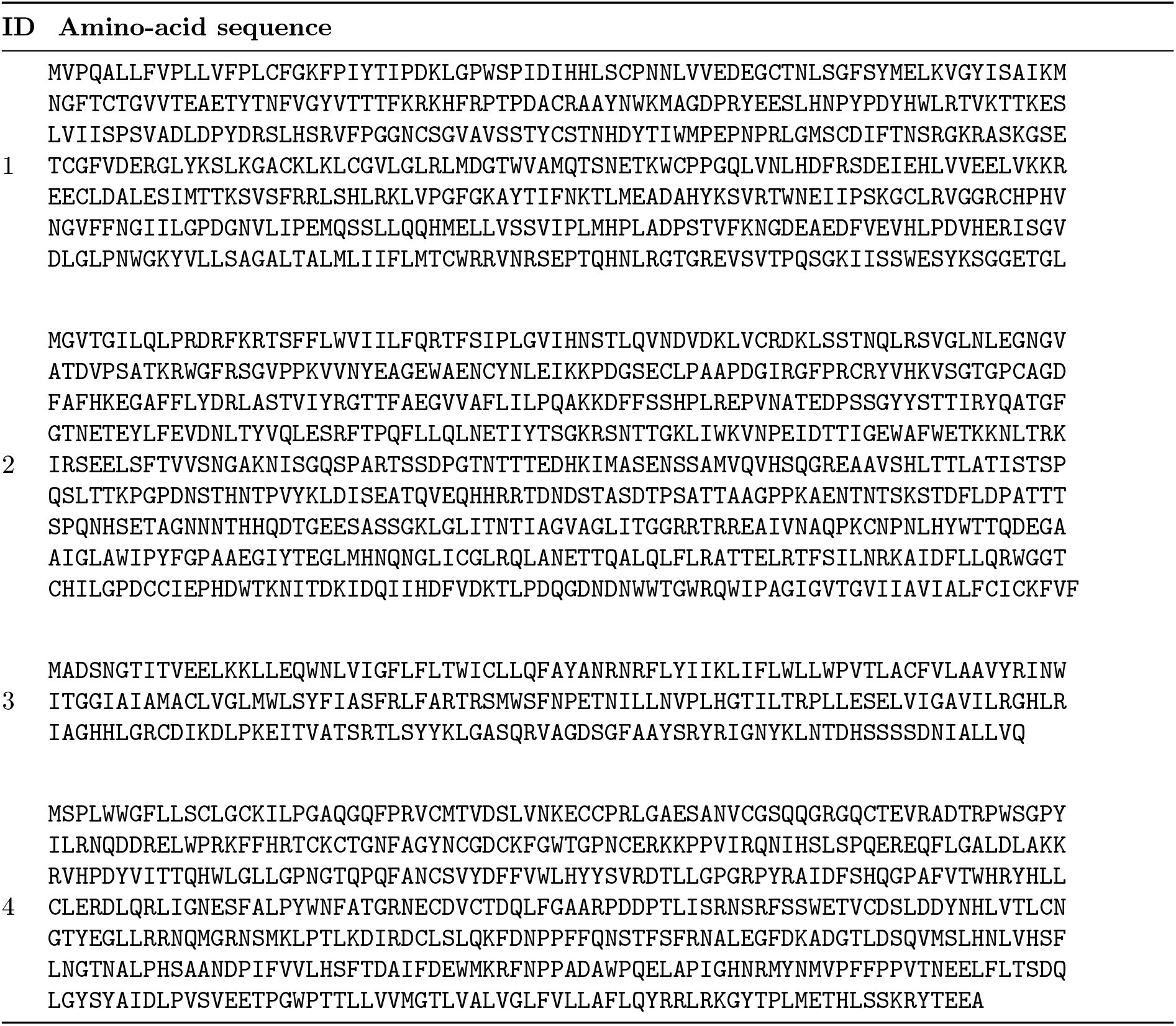
Amino-Acid sequences for protein panel used for CDS design and optimization.

**Figure S3:**
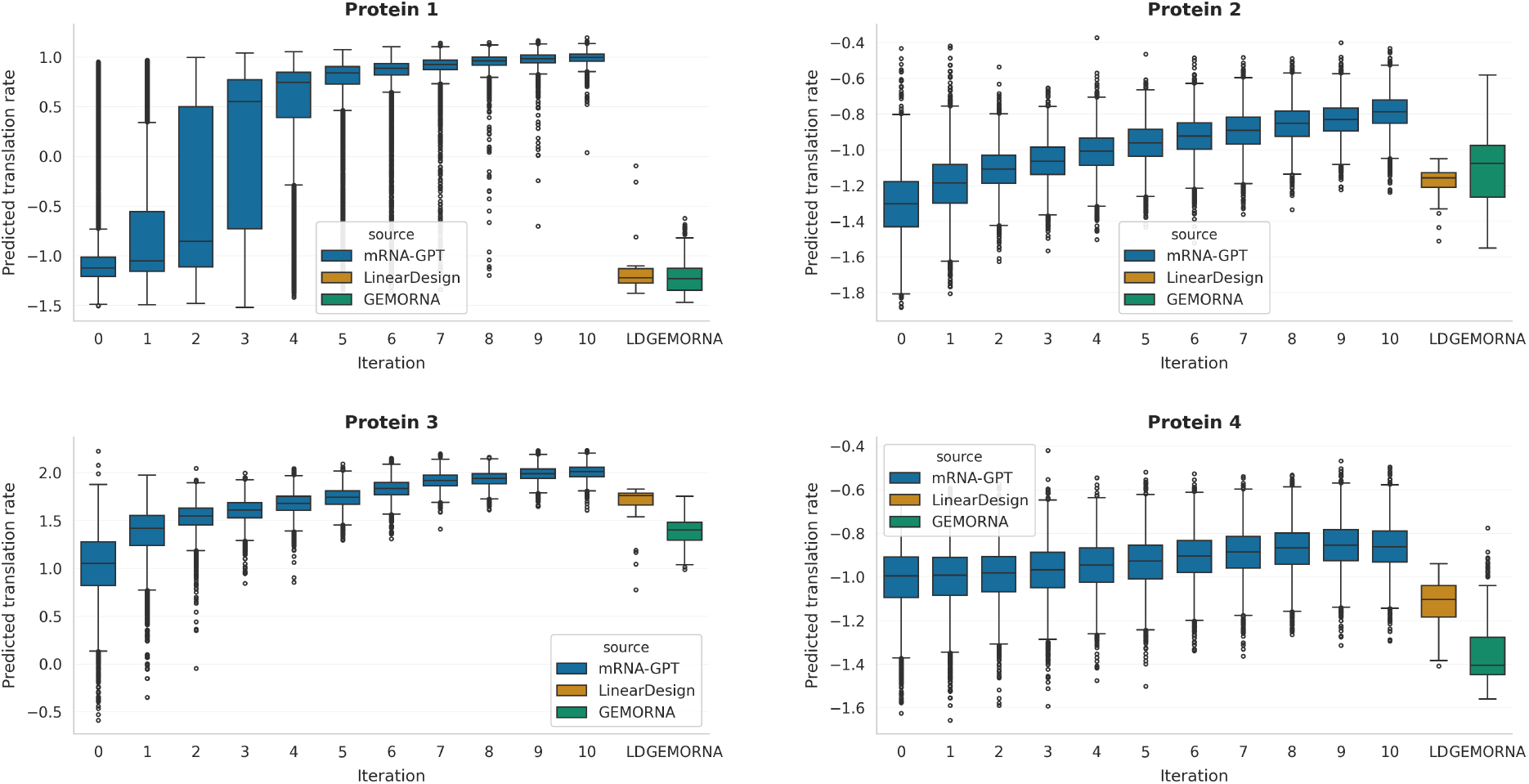
Oracle-predicted translation rate for CDS-only optimization across all four proteins. Each panel reports the distribution of mRNA-LM oracle scores for 6000 *CDS-only* sequences generated by mRNA-GPT per optimization round (*t*=0–10; blue boxplots), together with LinearDesign (“LD”, orange) and GEMORNA (green) CDS scored by the same oracle. Across all targets, the median and upper quantiles of mRNA-GPT steadily improve with iteration and low-scoring outliers are progressively suppressed, indicating batch-wide gains rather than improvements restricted to a few top sequences. For all proteins, the final mRNA-GPT iterations clearly outperform both baselines.

**Figure S4:**
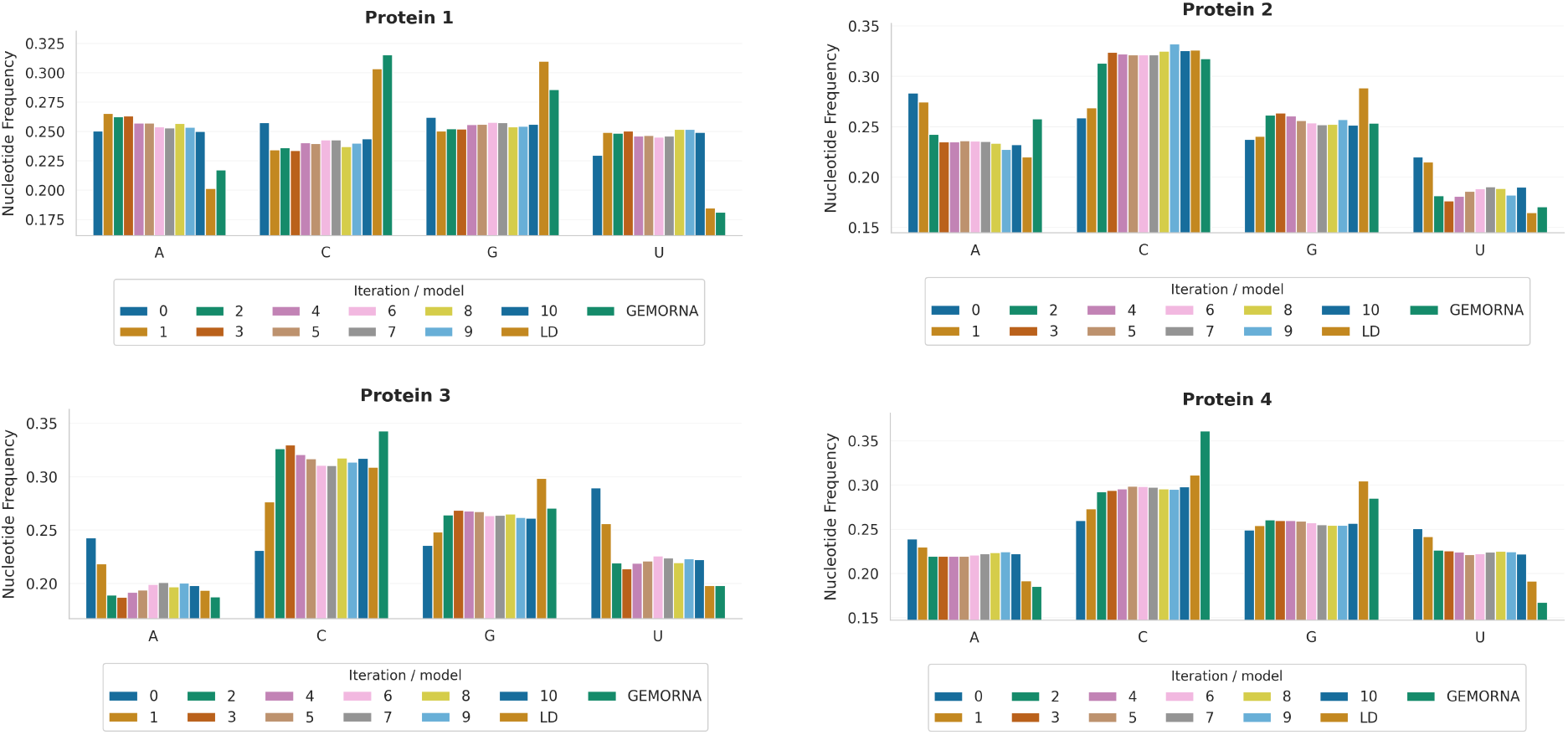
Nucleotide composition of CDS-only sequences across proteins and optimization rounds. Mean nucleotide frequencies (A, C, G, U) in the *coding sequences* optimized by mRNA-GPT are shown for each iteration (*t*=0–10; colored bars) and for the LinearDesign and GEMORNA CDS baselines, separately for the four target proteins. Across proteins, iterative CDS optimization increases GC content (higher C/G fractions with concomitant decreases in A/U) and converges toward the GC-rich regime explored by codon-design baselines, while preserving moderate, protein-specific variability in CDS composition.

**Figure S5:**
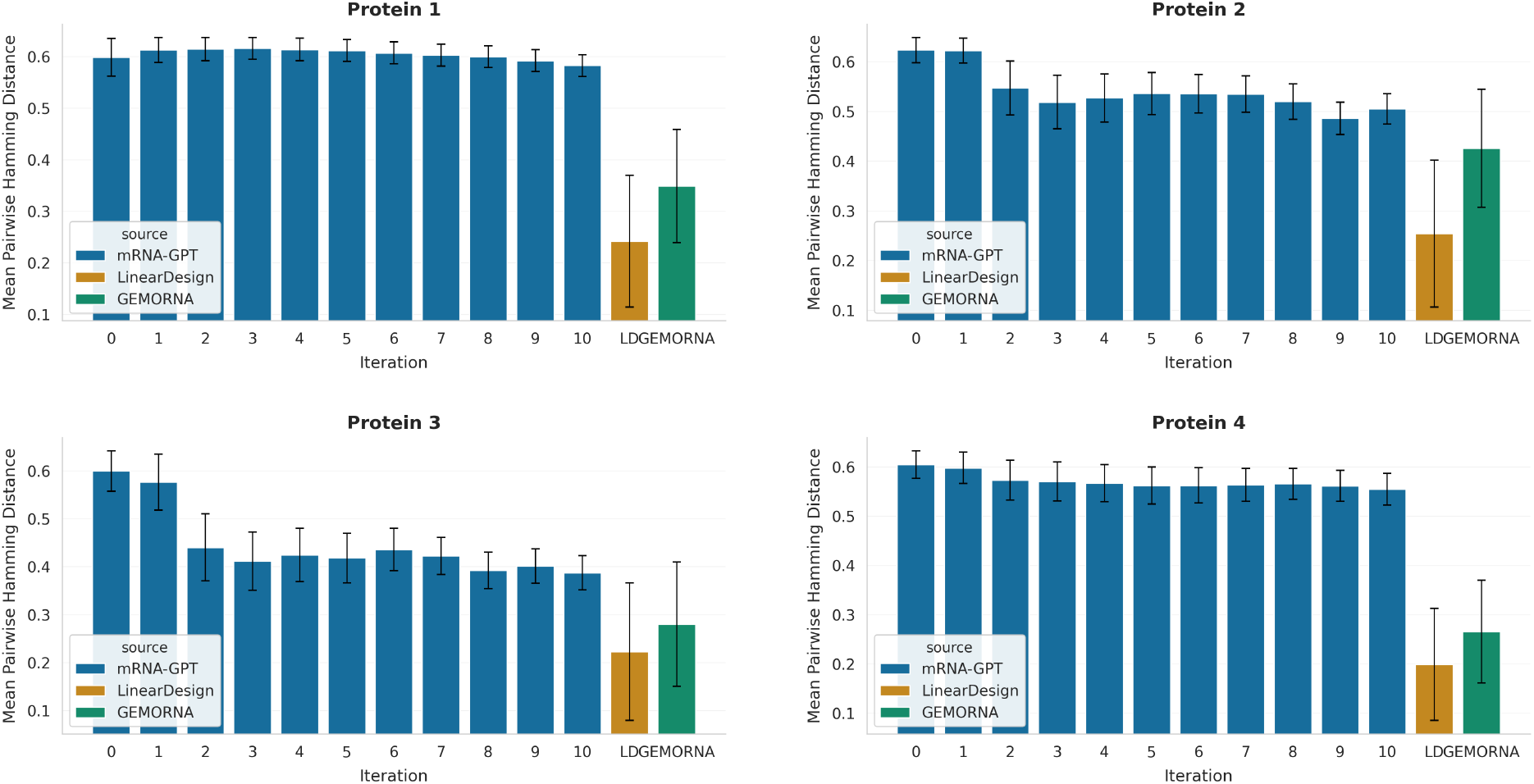
Per-protein codon-level diversity for CDS-only optimization (4 proteins). For each protein, bars report the mean pairwise codon-level Hamming distance between *CDS-only* sequences within a batch, with error bars denoting the corresponding standard deviation. Across all proteins, mRNA-GPT maintains a high degree of intra-batch diversity throughout CDS optimization. In contrast, LinearDesign and GEMORNA produce markedly less diverse CDS sets, indicating that their solutions concentrate in a narrower region of sequence space. These results show that mRNA-GPT improves CDS quality without collapsing to a small number of near-identical coding sequences.

**Figure S6:**
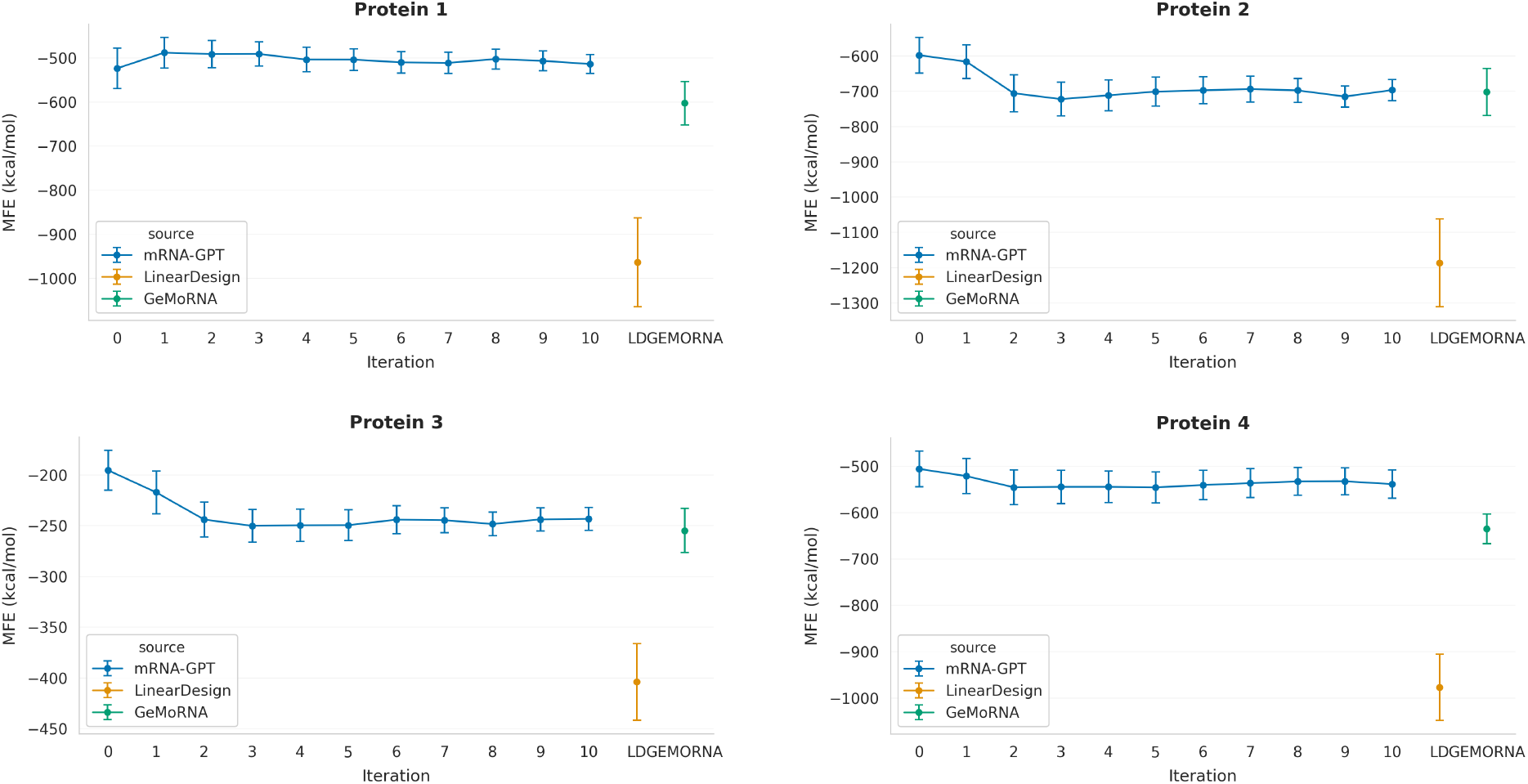
Per-protein minimum free energy (MFE) of optimized CDS-only ensembles. For each protein, blue points trace the mean predicted minimum free energy of mRNA-GPT-generated *CDS-only* sequences as a function of optimization iteration (*t*=0 to *t*=10); vertical bars denote one standard deviation over the corresponding sequence batch. Green and orange markers on the right-hand side summarize the mean ± s.d. MFE of GEMORNA and LinearDesign CDS, respectively, evaluated with the same RNAfold pipeline. Across all four proteins, mRNA-GPT operates in an intermediate MFE regime for CDS-only constructs, remaining systematically less structured (less negative MFE) than LinearDesign while typically matching or modestly improving on GEMORNA, indicating that CDS optimization does not collapse onto overly stable, potentially translation-inhibitory secondary structures.

**Figure S7:**
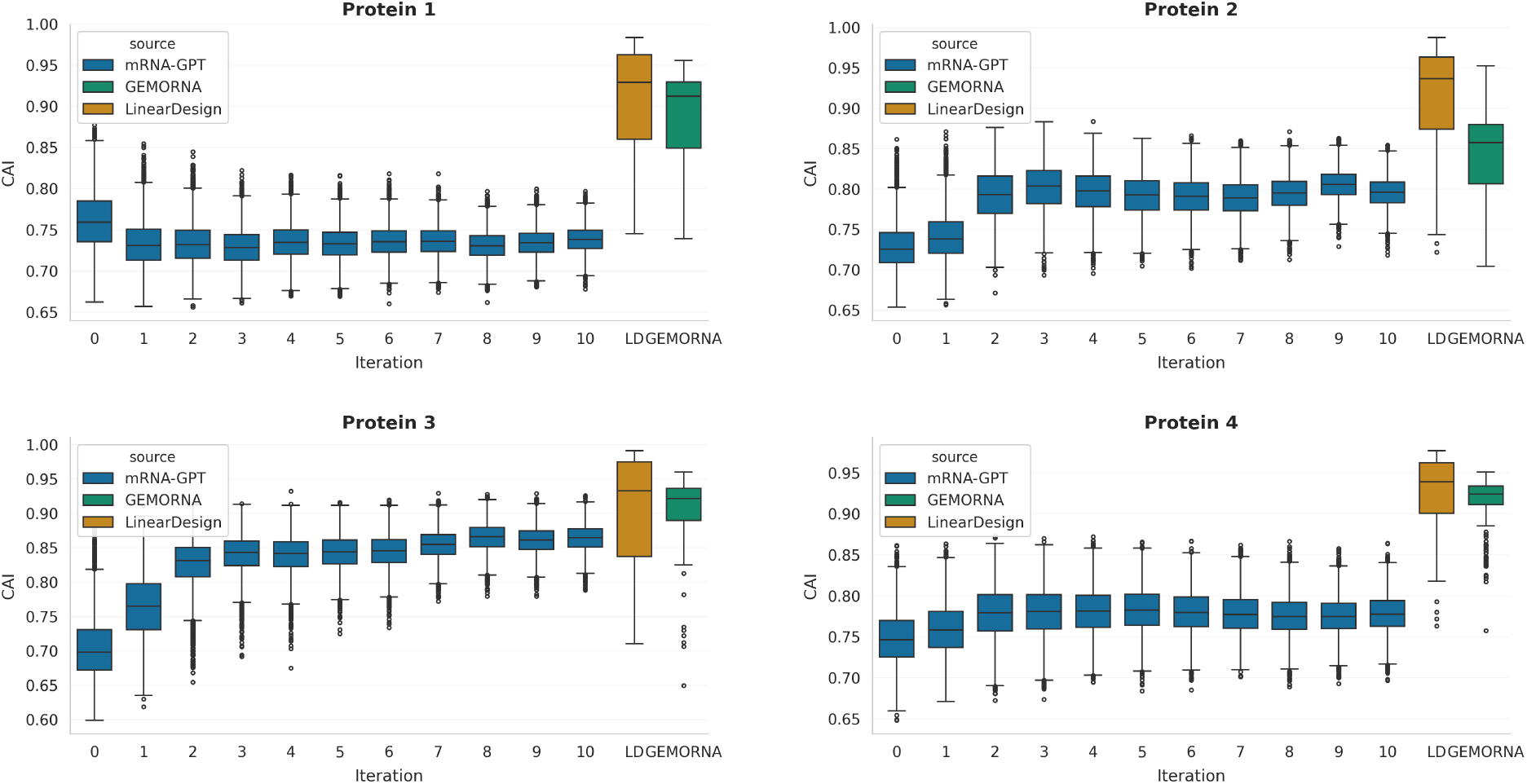
Per-protein Codon Adaptation Index (CAI) for CDS-only optimization over eleven rounds. For each protein, boxplots report the distribution of CAI scores for the mRNA-GPT– generated *CDS-only coding sequences* from the pre-trained model (*t*=0) to iteration *t*=10, with LinearDesign and GEMORNA CDS baselines plotted as separate groups on the right. Across proteins 2, 3 and 4, early optimization rounds (*t* ≤ 3) yield a pronounced upward shift and narrowing of the CAI distribution, after which median values plateau and only marginal additional gains are observed. LinearDesign and GEMORNA typically achieve higher absolute CAI values on CDS, but mRNA-GPT closes most of the gap while preserving broad sequence variability (see codon-diversity analysis) for protein 3 for instance.

**Figure S8:**
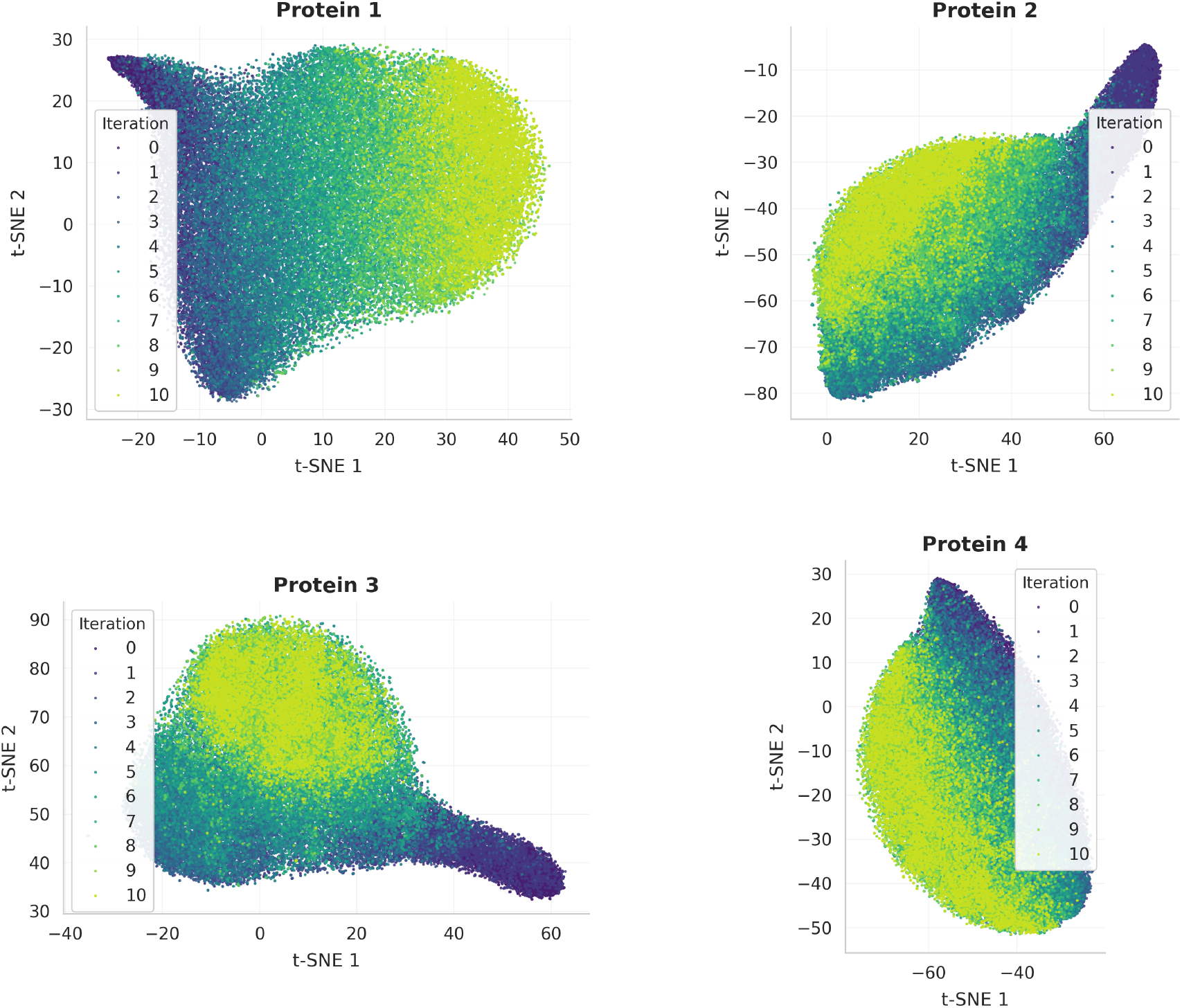
Per-protein t-SNE of CDS-only codon-usage profiles across optimization iterations. Two-dimensional t-SNE embeddings of 64-dimensional codon-usage vectors computed on the *CDS-only* portion of mRNA-GPT-generated sequences for Proteins 1–4. Each point corresponds to one CDS and is colored by optimization iteration (*t*=0–10). Across all targets, later iterations drift smoothly over the manifold explored by the pre-trained model while continuing to occupy a broad region of codonusage space, consistent with substantial optimization of the CDS-level oracle score without collapse onto a single codon pattern.

**Figure S9:**
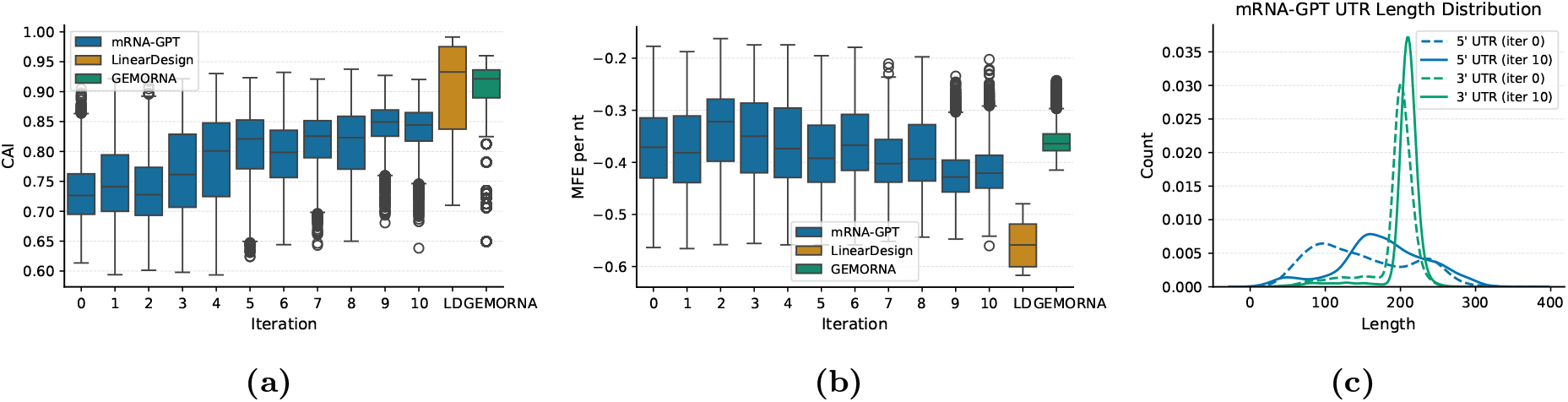
Distribution of CAI (a) and MFE (b) for mRNA-GPT sequences across optimization iterations, compared with GEMORNA and LinearDesign baselines. (c) UTR sequence length distributions.

